# What’s in it for the dogs? Assessing the outcomes of a prison-based dog training program from an animal behavior and welfare perspective

**DOI:** 10.1101/2024.10.15.618160

**Authors:** Parizad Baria-Unwalla, Gabriela Munhoz Morello, Maria Queiroz, I. Anna S. Olsson, Ana Catarina Vieira de Castro

## Abstract

Prison-Based Dog Training Programs (PBDTPs) are gaining popularity across the world for their benefits to inmates in terms of mental and emotional health, and reduced recidivism. However, research on the implications for the dogs is limited. The aim of the present study was to assess the outcomes of a PBDTP from an animal behavior and welfare perspective. Shelter dogs (N=42) were transported to and from the prisons twice a week where they received training from inmates for a total duration of 12 weeks. Dogs were tested for potential improvements in socialization and handling skills and basic training skills using standard tests (Temperament Test and Basic Education Test). Dog welfare was assessed using behavioral, physiological, and cognitive measures, namely stress-related behaviors and overall behavior states during training sessions, salivary cortisol levels, and cognitive bias tests. Results showed that participating in the PBDTP improved the dogs’ sociability towards humans and increased playful behavior, and improved basic training skills including not jumping on people, walking on a leash without pulling, responding to commands (sit, down, and staying in place) and staying calm when separated from the handler. Furthermore, behaviors indicative of stress were generally rare during training sessions and no impact of the PBDTP was found on the levels of salivary cortisol nor on the dogs’ affective states (as measured with cognitive bias tests). In conclusion, the present study suggests that PBDPTs are beneficial for dogs, with animals showing no indicators of compromised welfare while displaying improved behavior skills, which will likely facilitate their future rehoming.

## 1. Introduction

Correctional facilities across the world, including in USA [1], UK [2], Australia [3], Netherlands [4], and Japan [5] use different forms of prison-based dog programs to improve inmates’ social and emotional functioning and to reduce recidivism [6] [7] [8]. Dog training programs are one such form, along with other programs, including therapeutic interventions such as Animal Assisted Therapy.

Prison-based dog programs are based on the understanding that human-animal interactions bring psychosocial benefits including reduced stress, increased social support and emotional well-being [1]. The beneficial outcomes of Prison-Based Dog Training Programs (PBDTPs) on human participants (primarily inmates, but also staff at the correctional facility and the community at large) include increased offender self-control [9] [10] [11], increased self-esteem and self-efficacy [12] [13], increased empathy for dogs and humans [9] [10], increased emotional intelligence, and increased employability [1].

The popularity of PBDTPs has been on the rise worldwide in recent years which may be in part due to the positive media attention they receive and the public appeal of connecting shelter dogs and prison inmates, two vulnerable groups that need social contact [14] [15]. In the United States alone, over 330 correctional facilities have implemented some form of PBDTPs [16]. Research into PBDTPs has nearly exclusively focused on the impact on human participants [1], whereas there is hardly any data on the effect on dog health and welfare [17].

Dog training programs are potentially beneficial to dogs in several ways. Shelter dogs have higher chronic cortisol levels than owned dogs [18] [19], which is an indicator of higher stress levels in shelter dogs. However, interactions with humans at shelters, including walking and play, can reduce cortisol levels in dogs [20] [21], highlighting the potential benefits to dogs of positive human interactions. Many PBDTPs bring in dogs from shelters [1], which can potentially reduce their stress levels. PBDTPs may also create indirect benefits for the participating dogs. The training that these dogs receive may make them more attractive to potential adopters, and potentially facilitate their integration in a new family. Additionally, PBDTPs impart useful skills to dogs who sometimes go on to become service or working dogs and might have been euthanized otherwise [22]. However, participating in the training program also implies potentially stressful experiences, including change of environment, transport and interactions with unfamiliar and initially unexperienced handlers. Failing to consider the welfare of dogs in PBDTPs may put participants, facilitators, and dogs at risk [23].

There has been almost no research into how PBDTPs affect dogs, with a consequent lack of evidence regarding efficacy and welfare impact of these programs [23]. Hennessy et al [24] found that shelter dogs participating in a 3-week PBDTP exhibited increased training performance and decreased behavioral arousal in a novel situation from before to after the program as compared to control dogs that remained at the shelter. However, no differences were found between groups in physiological measures, with plasma cortisol levels remaining constant and pituitary hormone (ACHT) increasing from before to after the program. More recently, Leonardi et al [25] found that, after participation in a PBDTP, shelter dogs improved their performance in training tasks, showed increased levels of relaxed behaviors in their home kennels and received enhanced subjective ratings of behavior by shelter staff. Taken together, the results of these two studies suggest that PBDTPs improve dog training performance and do not impair dog welfare, with Leonardi et al [25] indeed suggesting a positive effect of participation in the program on the welfare of the animals.

While the results of these studies are promising, they are limited: Hennessy et al [24] only used physiological measures of stress (which provide information on the arousal but not on the valence of the animals’ emotions, e.g., [26]) and the main welfare measure used by Leonardi et al [25] comprised subjective scoring by non-blind raters. In addition, it is important to assess how participation in a PBDTP affects dogs both inside and outside of the training environment (i.e. the prisons).

Therefore, the aim of the present study was to perform a comprehensive evaluation of the effects of PBDTPs on dog behavioral prowess and welfare. To that end, we analyzed several indicators of dog socialization and handling and basic training before and after participation in a PBDTP and combined physiological, behavioral and cognitive measures to assess dog welfare before, during and after the program. Given the rising popularity of PBDTPs, systematic research on the topic is of critical importance.

## 2. Animals, Materials and Methods

### 2.1. Ethics Statement

Approvals were obtained from the i3S Committee for Ethics and Responsible Conduct in Research (N9/CECRI/2022) and from the i3S Animal Welfare and Ethics Review Body (internal reference 2021-29). All the involvement of the inmates and the dogs in the PBDTP was under the management and the responsibility of DTC Social (see below on ‘The Prison-Based Dog Training Program’). No identifying information was collected or recorded about inmates in the program for the purposes of the present study, and all results are presented without any disclosure of personal information. During training sessions at the prisons, datasheets were used to code dog behavior. These bore no information regarding the identity of the inmates training each dog and no visual recordings were made of sessions in the prisons. All collected data was coded on MS Excel and stored in encrypted folders on two different hard disks, exclusively for research use. Video recordings from the Temperament Tests (TTs), Basic Education Tests (BETs) and cognitive bias tests (described below) only featured the authors of the present study and the participating dogs.

No invasive or stressful tests were conducted with the animals: saliva sampling was non-invasive and implied only brief low-stress handling, behavior assessments during training sessions were purely observational and the behavior tests only implied tasks that are part of regular dog-human interactions for family dogs.

The researchers worked under the close supervision of the DTC Social team, who were responsible for ensuring the necessary health and safety protocols.

### 2.2. The Prison-Based Dog Training Project

Pelos2 is a PBDTP conducted by DTC Social, a Portuguese NGO that works with dogs, and funded by Portugal Inovação Social, a public initiative to promote social innovation in Portugal. The program, based on the training of shelter dogs by prison inmates, has a twofold aim: 1) imparting inmates with occupational skills and reducing the impact of incarceration on their mental health, and 2) granting dogs with educational skills and attempt at promoting their successful rehoming.

The research team at i3S (i.e., the authors of this paper) collaborated with DTC Social to collect research data on dogs participating in the program in three male prisons in the north of the country: Santa Cruz do Bispo, Vale do Sousa, and Paços de Ferreira

### 2.3. Participants

Expecting high variability in the sample, even relatively large differences might yield small effect sizes. Hence, we powered this experiment to detect a Cohen’s f=.25. Power analysis (G*Power) for a repeated-measures design deemed N=34 sufficient (α=.05, β=0.8). Forty-five dogs were recruited for the present study and equally allocated by DTC Social to three male prisons: Paços de Ferreira (Prison 1), Vale do Sousa (Prison 2), and Santa Cruz do Bispo (Prison 3). During the course of the program, however, three dogs dropped out (two due to health issues and one due to behavior management issues at the prison); hence we ended up with a final sample of 42 animals.

The participating dogs were selected by DTC Social from the Plataforma de Acolhimento e Tratamento Animal (PATA), a municipal companion animal shelter in the Porto region. Selected dogs were between 1-8 years of age, healthy, without visible signs of aggression or extreme fear (as determined by DTC trainers in a brief interaction with the animals), and had been at the shelter for at least three weeks before the PBDTP commenced. While it was attempted to recruit an equal number of male and female dogs, the predefined eligibility criteria and availability of dogs at PATA did not allow this, hence we ended up with 12 female and 33 male dogs in the program. Detailed demographics are depicted in Appendix 1.

For the duration of the program, these dogs were not available for immediate adoption (i.e., potential adopters were able to place the dogs on hold but could only take them home once their participation in the program was complete). The information regarding their participation in the training program was displayed on a sign at their home enclosures at PATA.

### 2.4. Experimental Design and Data Collection

#### 2.4.1. Training Program

The basic training skills imparted to the dogs included sit, down, stay, come when called, loose leash walking, and not jumping on people. The socialization and handling skills included sociability with dogs and humans, handling for hygiene, grooming and veterinary procedures, resource guarding, confidence, and independence. Only reward-based methods (positive reinforcement training) were used. An overview of the training plan followed by the DTC Team is presented in Appendix 2.

The dogs were picked up from PATA at 9 am and returned at 4 pm, from Monday to Thursday. Each dog was allocated to one prison, and transported there two days a week (on alternate days). On these days, they participated in three different sessions, each with a different inmate. Each session was 45 minutes in duration, including walking, playtime, and 15 minutes of dedicated training. In between training sessions, the dogs remained resting in their crates in the van for 15 to 90 minutes (15 minutes between subsequent sessions, up to 90 minutes during lunch break). The vans were equipped with ventilators and were always parked in shaded areas, with the doors at the back and side left ajar enough that there was enough ventilation at all times. On Friday, Saturday, and Sunday, all dogs remained at PATA. At PATA, the dogs were kept in individual kennels and were not routinely handled beyond the basic human interactions around feeding and kennel cleaning. They were fed between 8:00 and 8:30 am every day at the shelter and had water available throughout the day, including during the training sessions at the prisons.

The program lasted 12 weeks each, for four different batches of dogs. The first two batches included 6 dogs each, the third batch included 17 dogs and the fourth batch 16 dogs (after accounting for dropouts).

#### 2.4.2. Testing

A detailed schedule for the testing timeline is provided in Figure 1.

**Figure 1.**
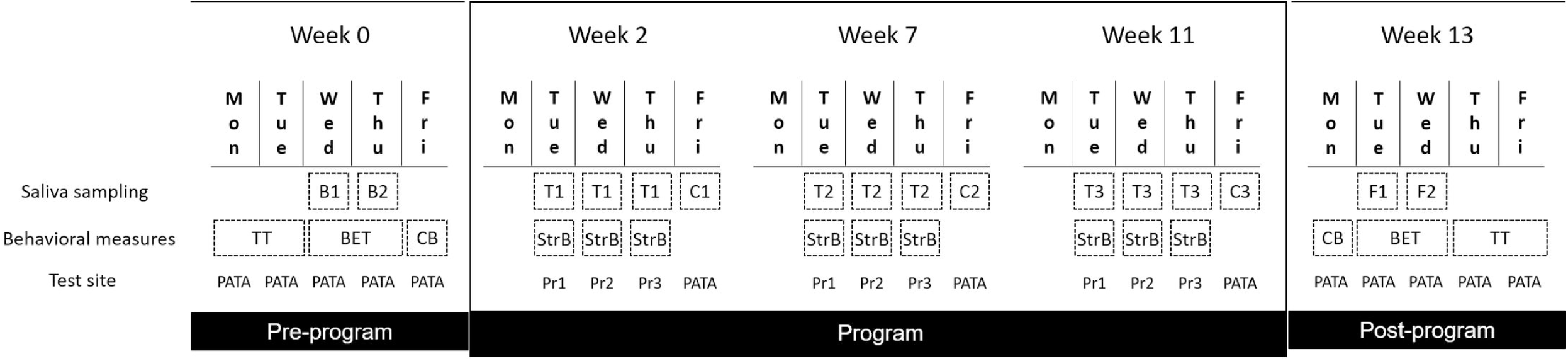
Study timeline. Dog socialization and handling and basic training indicators were assessed with TT - Temperament Test and BET – Basic Education Test, respectively, and dog welfare with salivary cortisol concentration (B1, B2, C1-C3, T1-T3, F1, F2), stress-related behaviors (StrB) and cognitive bias tasks (CB). Dogs were tested with TT, BET and CB before (pre-program, week 0) and after participation in the PBDTP (post-program, week 13). TTs and BETs were conducted over two days at each stage (half of the subjects on each day). Stress-related behaviors were assessed during PBDTP training sessions at the prisons (Pr1- Prison 1; Pr2 – Prison 2; Pr3 – Prison 3) at three time-points: week 2, week 7, and week 11. Saliva samples T1-T3 (T; training) were collected on these same days at the prisons; and samples C1-C3 (C, control) were collected at PATA shelter at the same time-points (week 2, week 7, week 11) but on a different day, when the dogs were not transported to the prisons and remained at the shelter. B1 and B2 are the pre-program baseline samples and F1 and F2 the post-program final samples collected at PATA on the same days the BETs were conducted.

For evaluating the changes in socialization and handling and basic training skills, the dogs were tested at two time-points: one in the week just before commencing the PBTDP (pre-program, week 0) and one in the week just after completion of the PBDTP (post- program, week 13). They were tested for basic training skills using the Basic Education Test (BET), adapted from the Canine Good Citizen Test by the American Kennel Club (AKC) [27], and for socialization and handling skills using the Temperament Test (TT), developed specifically for use in shelter contexts by Valsecchi et al [28] and further refined by Barnard et al [29]. Both testing protocols (described in detail in Appendices 3 and 4) have been previously piloted by us [30]. For standardization purposes, the handler in the BETs was played by Maria Queiroz (MQ) for both stages. For the TT, Ana Catarina Vieira de Castro (ACVC) and MQ took turns playing the handler in the pre- and post-program assessments (MQ in the pre- and ACVC in the post-program assessments), so that the dogs were always tested by a relatively unfamiliar person. The tests were scored live using established scoring systems (see Appendices 3 and 4) and were also video recorded to allow further analysis. ACVC live scored the dogs in the BETs and in the pre-program TTs. MQ live scored the post-program TTs.

For evaluating dog welfare during the participation in the PBDTP, behavioral and physiological measures were used. The behavior of the dogs during the first training session of the day at three time-points of the PBDTP (weeks 2, 7, and 11) was coded live using previously validated ethograms (Appendices 5 and 6) by ACVC and Parizad Baria-Unwalla (PB). Each researcher coded the behavior of three different dogs, following each dog for five minutes. They used continuous sampling for coding stress-related behaviors, including lip- licking, yawning, moving away, crouches, body-turns, and body-shakes (as per definitions in Appendix 5), and instantaneous scan sampling every 60 seconds for coding overall behavior states (Tense, Low, Relaxed, Excited, as defined in Appendix 6). The ethograms were based on studies of dog welfare in a training context [31] [32], and include commonly observed behaviors during training and especially those most prone to show differences in stressful vs non-stressful conditions. The two researchers coding dog behavior during the training sessions trained and calibrated themselves for inter-observer reliability from previously recorded dog training videos from a different project, until a high level of agreement was achieved (>90% for the frequency of stress-related behaviors; >70% for overall behavior states). For stress-related behaviors, the reliability was checked using intra-class correlation and for overall behavior states, it was checked using Cohen’s Kappa statistic.

Salivary cortisol concentration was used as a physiological measure of welfare, in conjunction with the behavioral measures [31] [32]. Samples of dogs’ saliva were collected with cotton buds (Salivette^®^). One of the researchers used a cotton bud and moved it in gentle circular motions on the inside of the dogs’ cheeks, while stroking and praising the dog. Before the program commenced, two baseline samples (B1 and B2) were collected at the shelter between 11:00 and 15:00 in two different days (the same days on which the BETs were performed, and immediately after taking the dogs out of their home pens for testing, see Figure 1). Once the PBDTP started, samples were collected between 12:30 and 14:30 on training days at the prisons during weeks 2, 7, and 11 (training samples; T1, T2 and T3). The dogs had undergone training sessions in the morning and had rested in the crates/vans for at least one hour prior to saliva collection. Samples were also collected during the same hour range at PATA on Fridays of the same weeks, when the dogs were at the shelter all day (control samples; C1, C2 and C3). After the program ended, two other samples (F1 and F2) were collected at the shelter between 11:00 and 15:00 in two different days (once again, the same days on which the BETs were performed, see Figure 1). The goal of collecting B samples was to allow investigating (i) whether C samples could be used as true controls for T samples (i.e., whether the cortisol levels of the animals when at the shelter would not change with the participation in the PBDTP) and (ii) together with F samples, examining whether participating in the PBDTP would affect dogs’ cortisol levels at the shelter outside the program period. All samples were labelled with the dog ID, date, time, stored in individual plastic tubes on ice, and moved as soon as possible to a -20 Celsius freezer at i3s. At the end of data collection for each batch, the samples were sent to an external laboratory (DNAtech) where they were tested for cortisol concentration with ELISA.

Cognitive bias tests were also carried out before (week 0) and after the PBDTP (week 13) to evaluate the effects of the program in the affective state of the dogs. Dogs who were unmanageable on the leash during the Temperament Test were excluded from selection for this test, as well as dogs who showed no interest in food. The cognitive bias tests were conducted in an indoor room at PATA. The dogs learned a discrimination between two food bowl locations (positive, which contained food, and negative, which was empty) and were then tested with an ambiguous (empty) food bowl, equally spaced between the two training locations. The time (in seconds) dogs took to reach the food bowls was scored live using a stopwatch. The entire protocol is detailed in Appendix 9.

Due to logistical constraints, the TT and BET tests were conducted for all batches (1- 4), whereas smaller subsets of dogs were tested for cortisol levels, stress-related behaviors and cognitive bias. A total of 42 dogs underwent TT testing during both the pre- and post-program assessments. However, two dogs (marked with * in Appendix 1) refused to walk on a leash during the pre-program BET, and thus could not be tested at that stage. Consequently, 40 dogs were tested in both the pre- and post-program BET. Moreover, four dogs (marked with ** in Appendix 1) did not allow handling for saliva collection and were excluded from cortisol sampling to avoid causing unnecessary stress. C and T samples were then collected for a total of 34 dogs (from batches 2-4) and samples B and F for a sub-sample of 13 dogs (batch 4). Stress-related behaviors were coded for 33 animals (from batches 3 and 4) and cognitive bias testing was conducted for a sub-sample of 16 dogs (from batches 3 and 4). Sub-samples were balanced for prison and sex.

The sequence in which the TT, BET, and cognitive bias tests were administered during the pre-program assessment was determined with safety and standardization in mind. Since the dogs were unfamiliar to the research team, conducting the TTs first allowed for an evaluation of their behavior around humans and their response to handling in a relatively safe setting, before progressing to the more intrusive handling required for BET and cognitive bias tests. Cognitive bias tests were not performed on the first two batches, which were then tested with the BET immediately following the TT. To maintain consistency across batches, the cognitive bias test was conducted as the final assessment in the test battery. For the post- program assessment, we changed the order of the tests to 1) cognitive bias, 2) BET, and 3) TT, in order to prioritize those with the least potential to influence the outcomes of subsequent tests (see Figure 1).

### 2.5. Data analysis

In addition to the live scoring of the dogs’ performance in BETs and TTs, the tests were also scored from video recordings (using the same scoring systems - Appendices 3 and 4). The videos for each batch of dogs were analyzed by four observers, three blind to the stage (pre- vs post-program) and to the goals of the study, and one non-blind (assisted in data collection). Different blind observers (selected from a pool of eight) were allocated to different batches; the non-blind observer scored all batches (see Appendices 7a and 7b). Inter-observer reliability (for the five observers) was calculated for each exercise using intra-class correlation coefficient (ICC, see Appendices 8a and 8b). The live and video scorings were combined into a final score for each BET and TT item. The average scores from the video observers comprised 50% of the total weightage. The remaining 50% was allocated to live scores by MQ and ACVC, both with extensive experience with dogs (details in Appendices 7a and 7b).

The BET measured variables across seven different categories, with one score per category – ‘Approach’ (Item 1 - Accepting a friendly stranger), ‘Greet’ (Item 2 – Waiting politely for petting), ‘Leash’ (Item 3 – Walking on a loose leash), ‘Crowd’ (Item 4 – Walking through a crowd), ‘Commands’ (Item 5 – Sit, down and staying in place), ‘Recall’ (Item 6 – Coming when called) and ‘Separation’ (Item 7 – Supervised separation). A Total BET score was also computed by summing the scores of the different categories.

Although the TT was designed to measure variables across seven different categories (Behavior in Kennel, Human Sociability, Behavior on Leash, Cognitive Skills, Playfulness, Reactivity and Dog Sociability, see Appendix 4), only three categories i.e. Human Sociability, Dog Sociability, and Playfulness emerged as robust measures in previous studies [27] [28], therefore we limited the analysis to these same three categories. They were calculated as follows:

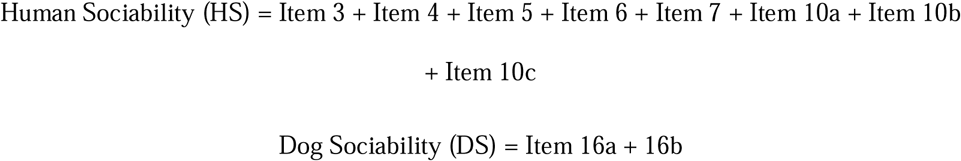

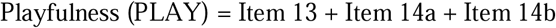

A Total TT score was also computed by summing the scores of all the items.

In the cognitive bias test, the dogs’ latencies to reach the bowl in the ambiguous location was adjusted for the latency to reach the training (positive and negative) locations (Mendl et al, 2010; see Appendix 9), as follows:

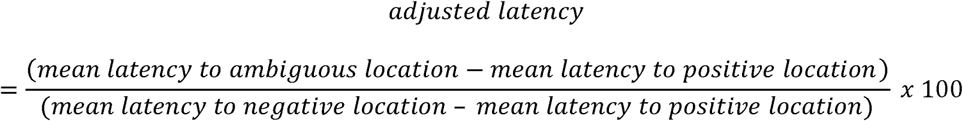

The frequency of stress-related behaviors and the frequency of scans in the different overall behavior states were input for analysis as recorded live by the researchers, and salivary cortisol concentrations (in ng/mL) as reported by the laboratory that ran the ELISAs (DNAtech).

#### 2.5.1. Statistical analysis

Prior to the main modeling, we tested the adequacy of using cortisol levels obtained on non-training days at the shelter in the course of the PBDTP (C1 to C3) as control levels (covariates) for cortisol levels obtained on training days at the prisons (T1 to T3). To this end, we tested if they differed from baseline cortisol levels at the shelter before the program started (B1 and B2). We also tested if dogs’ cortisol levels at the shelter changed from before to after the PBDTP, by comparing cortisol levels at the shelter before the program started (B1 and B2) with cortisol levels at the shelter after the program ended (F1 and F2). Paired t-tests were performed on SAS University Edition® (SAS Institute Inc., Cary, NC, USA) to compare baseline cortisol levels measured during the week prior to the start of the PBDTP (average of B1 and B2) with C1 to C3, and with post-program cortisol levels measured during the week after the completion of the PBDTP (average of F1 and F2). Some saliva samples did not have enough volume to allow laboratory analysis, which led to missing values. When both B1 and B2 levels were available, average B was obtained and used for comparison; when one of the B readings was missing, the available B level was considered for the comparisons. The same approach was taken for F cortisol levels. As a result, paired t-tests were done in sample subsets of distinct number of dogs (see Appendix 10).

The response variables training cortisol concentration (T), as well as Total TT Score, Human Sociability, Dog Sociability, Playfulness, Total BET Score, BET categories ‘Approach’, ‘Greet’, ‘Leash’, ‘Commands’ and ‘Separation’, the number of occurrences of the stress-related behavior Lip-licking, the number of scans in the behavior states Tense, Excited, and Relaxed, and the adjusted latency to reach the ambiguous bowl in the cognitive bias task were modeled as functions of the independent variables time-point (3-level categorical fixed effect for stress-related behaviors and behavioral states/2-level categorical fixed effect for remaining response variables), prison (3-level categorical fixed effect), the interaction between these two variables, dog age (discrete numeric fixed effect), and weight score (discrete fixed effect). Information on dog weight was estimated from visual inspection and input in ranges of 5 kg by DTC team (see Appendix 1), but for purposes of statistical analyses we created a weight score from one to seven, where one was 5-10 kg, two was 10-15 kg, three was 15-20 kg, four was 20-25 kg, five was 25-30 kg, six was 30-35 kg, and seven was 40-45 kg.

An adapted bi-directional stepwise selection of variables was used, starting with the independent variable of interest (time-point), followed by prison, the interaction between these two variables, dog age (discrete fixed effect), and weight score. Variables reaching significance (i.e. p≤0.05) were maintained in the model. Sex and neuter status were also tested as independent variables, but always one at a time, due to the unbalanced nature of the dataset. The effect of neuter status in the response variables were only tested for male dogs, as there was only one intact female dog in the dataset. Control cortisol concentration (C) was used as a covariate in the model explaining the variation of training cortisol concentration (T) as a function of the aforementioned effects. Procedure GLIMMIX (for generalized linear mixed modelling) was used on SAS University Edition® (SAS Institute Inc., Cary, NC, USA), considering Dog ID as a random effect.

Multicollinearity was checked by regressing each independent variable on the others in a numerical format. When model residuals failed to reach normality, a more appropriate data distribution was considered in the models: log-normal for TT category Human Sociability, behavior state Excited, training cortisol concentration (T), and Poisson for the stress-related behavior Lip-licking, behavior states Relaxed and Tense, and adjusted latency to reach the ambiguous bowl. To allow for the log-normal distribution in the models, 0.1 was added to all data points of the response variables to eliminate all zeros of the datasets. Adjusted latency was normalized prior to the analysis, so that the maximum adjusted value was 1 and the minimum was 0, to reach normality of model residuals.

The remaining stress-related behaviors Body Shake, Body Turn, Crouch, Move Away, and Yawn, as well as the behavior state Low, had no occurrences in 64%, 77%, 83%, 85%, 88%, and 96% of the observations, respectively, rendering impossible to explain their occurrences as a function of the variables of interest. Therefore, the occurrences of these stress-related behaviors were dichotomized into “1” (having occurred at least once) and “0” (no occurrences), and Fisher’s exact test was used through Procedure FREQ on SAS University Edition® (SAS Institute Inc., Cary, NC, USA) to evaluate the contingency between the 2-level categorical response variable (“0” or “1”) and Time-point for each of the participating prisons.

BET categories ‘Crowd’ and ‘Recall’ were dichotomized into being low (≤ 2.92, ≤ 1.8, respectively, thresholds based on data distribution) or high (higher than thresholds), and Procedure GLIMMIX was used considering a binary data distribution, with a Logit link function, to model probability of score being high as a function of the aforementioned independent variables.

For all fitted models, least squares means were compared considering 95% confidence interval (i.e. significance at p<0.05). A Tukey-Kramer adjustment was applied to all multiple comparisons performed in this study.

## 3. Results

### 3.1. Dog sociability and education indicators

#### 3.1.1. Basic Education Test (BET)

Figure 2 depicts the average scores for the BET categories pre- and post-program. We found that post-program scores were significantly higher compared to pre-program scores for all but one category: ‘Approach’ [F_(1,39)_=10.53, p=0.002], ‘Greet’ [F_(1,39)_=12.25, p=0.002], ‘Leash’ [F_(1,39)_=5.39, p=0.025], ‘Crowd’ [F_(1,439_=4.36, p=0.044], ‘Commands’ [F_(1,39)_=54.63, p<0.001], and ‘Separation’. The only category for which there was no significant effect of stage was ‘Recall’ [F_(1,39)_=0.31, p=0.579]. Total BET score also increased significantly from before to after the training program [Pre-program (Mean±SEM): 13.03±0.30; Post-program (Mean±SEM): 15.39±0.40; F_(1,39)_=69.95, p<0.001].

**Figure 2.**
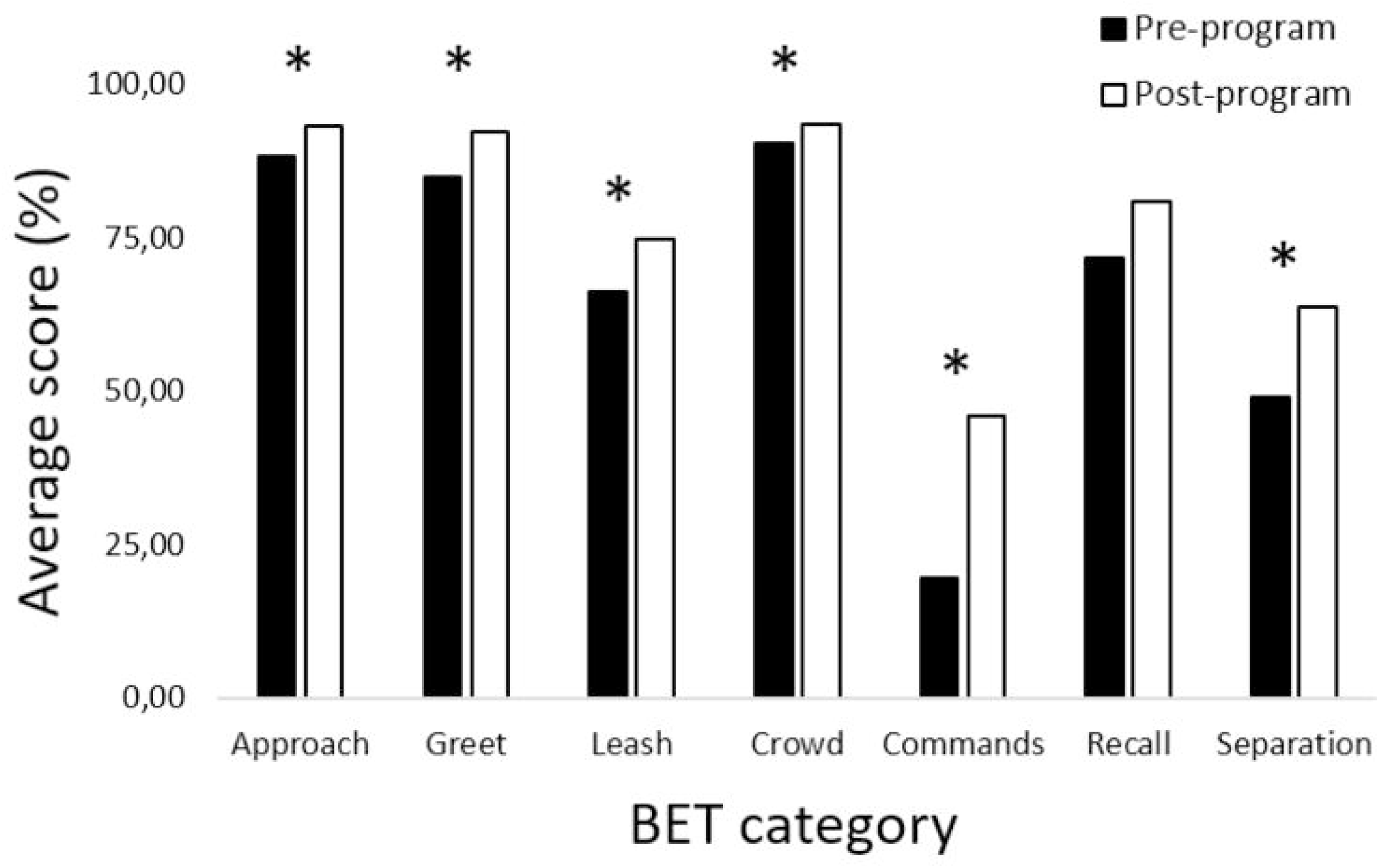
Average scores per category (expressed as a percentage of highest possible score) of the Basic Education Test, before (black bars) and after (white bars) the program (N=40). *stands for statistically significant differences (p<0.05).

#### 3.1.2. Temperament Test (TT)

Figure 3 depicts the average scores for the categories Human Sociability, Dog Sociability and Playfulness of the Temperament Test before and after the PBDTP. We found that the scores for Human Sociability and Playfulness were significantly higher after the PBDTP compared to before the PBDTP [Human Sociability: F_(1,41)_=8.24, p=0.006; Playfulness: F_(1,41)_=7.01, p=0.012], but we found no significant effect of stage (pre- vs post- program) on Dog Sociability [F_(1,39)_=0.13, p=0.724]. Results also showed that the Total TT score increased significantly from pre-program to post-program [Pre-program: 48.16±1.46; Post-program: 51.07±1.32; F_(1,41)_=16.45, p=0.0002]. Moreover, Playfulness was significantly affected by sex, with males exhibiting higher scores than females [F_(2,59)_=4.22, p=0.046].

**Figure 3.**
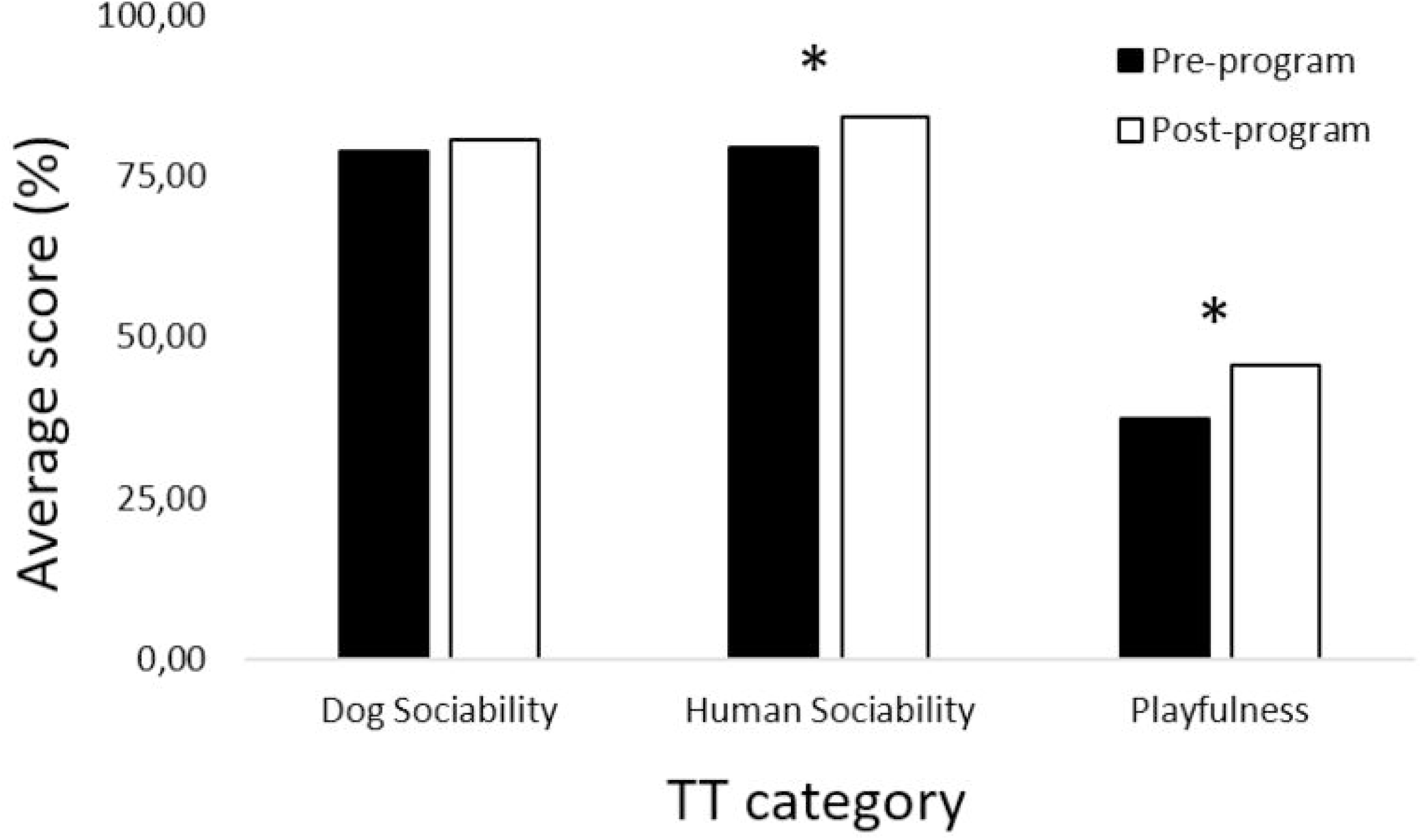
Average scores for the categories Human Sociability, Dog Sociability and Playfulness of the Temperament Test (expressed as a percentage of highest possible score), before (black bars) and after (white bars) the program (N=42). *stands for statistically significant differences (p<0.05).

### 3.2. Dog stress and welfare indicators

#### 3.2.1. Stress-related Behaviors

Lip-licking behavior was affected by time-point [F_(2,59)_=14.91, p<0.001], in that it was more frequent for the first time-point (Week 2) compared to the second (Week 7; t=4.25, p<0.001) and third (Week 11; t=4.89, p<0.001) time-points, as shown in Figure 4. No effects of time-point were found for the behaviors Yawn, Move Away, Body Turn, Crouch, and Body Shake, in any of the participating prisons (p>0.05). We also found that Lip-licking was affected by prison [F_(2,30)_=4.17, p=0.025], with dogs lip licking more frequently in Prison 2 compared to Prison 1 (t=-2.62, p=0.035).

**Figure 4.**
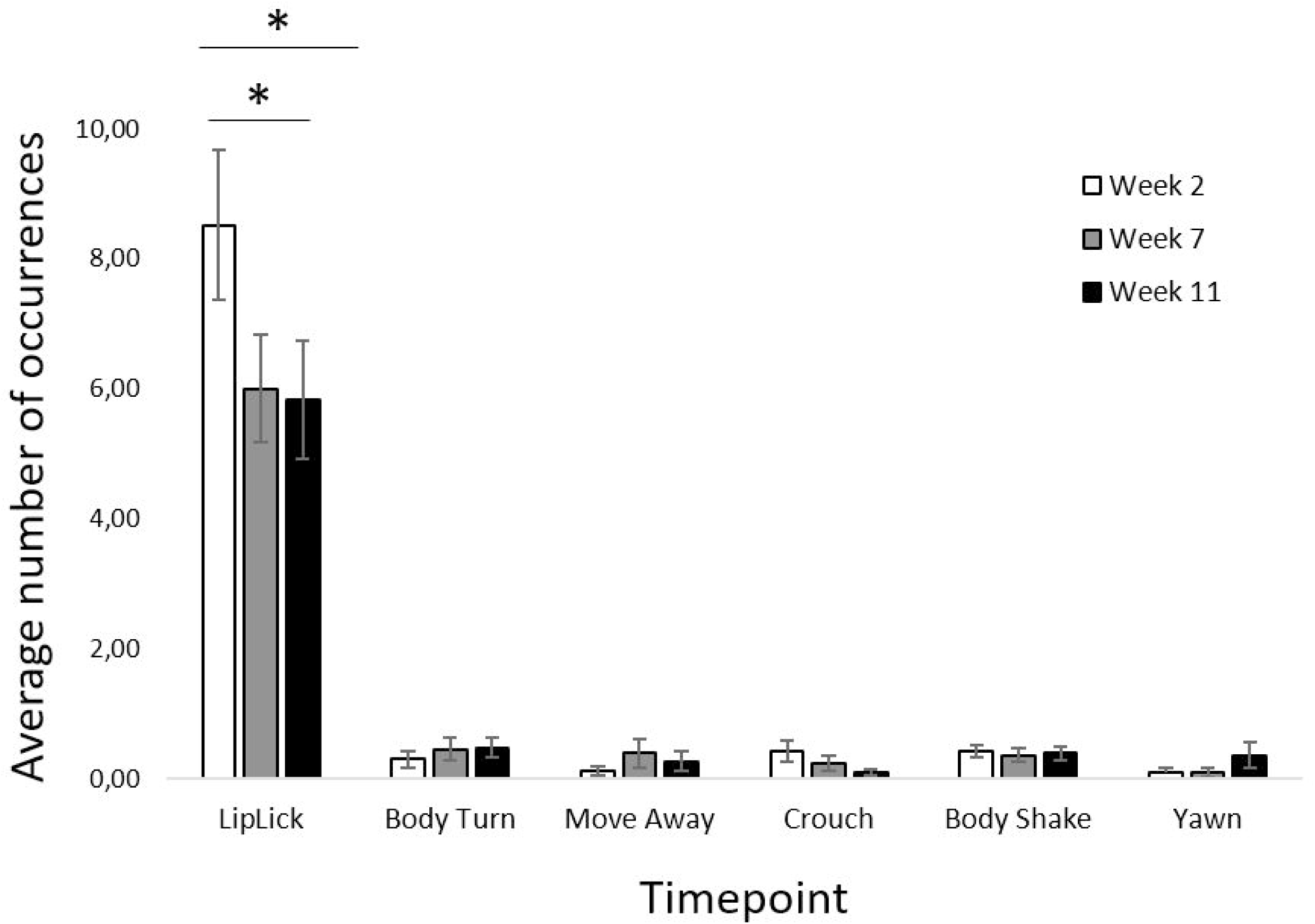
Average number of occurrences of stress-related behaviors during training sessions at the prisons across three time-points (Week 2, Week 7 and Week 11; N=33). Vertical bars show the SEM. *stands for statistically significant differences (p<0.05).

#### 3.2.2. Overall Behavior States

We found an effect of time-point on the frequency of scans Relaxed [F_(2,59)_=3.60, p=0.034] and Tense [F_(2,59)_=3.74, p=0.030]. Pairwise comparisons revealed that dogs were found more frequently in a Relaxed state and less frequently in a Tense state for the second time-point compared to the first time-point (Relaxed: t=-2.60, p=0.031; Tense: t=2.45,p=0.045, see Figure 5). No effect of time-point was found for the behavior states Excited [F_(2,55)_=0.10, p=0.902] and Low [F_(2,59)_=3.64, p=0.032]. We also found an effect of neuter status for the behavior state Tense on male dogs [F_(1,59)_=4.19, p=0.045]. This effect was not tested for females because there was only one intact female. A paired-samples t-test conducted for male dogs revealed that intact males were found less frequently in Tense states than neutered males (t=-2.05, p=0.045). Finally, we found an effect of weight score for the behavior state Relaxed [F_(1,59)_=8.25, p=0.006], with the frequency of scans increasing as weight score, thus dog weight, increased, and an effect of age on the behavior state Excited [F_(1,61)_=4.30, p=0.042], with the frequency of scans increasing as dog age decreased.

**Figure 5.**
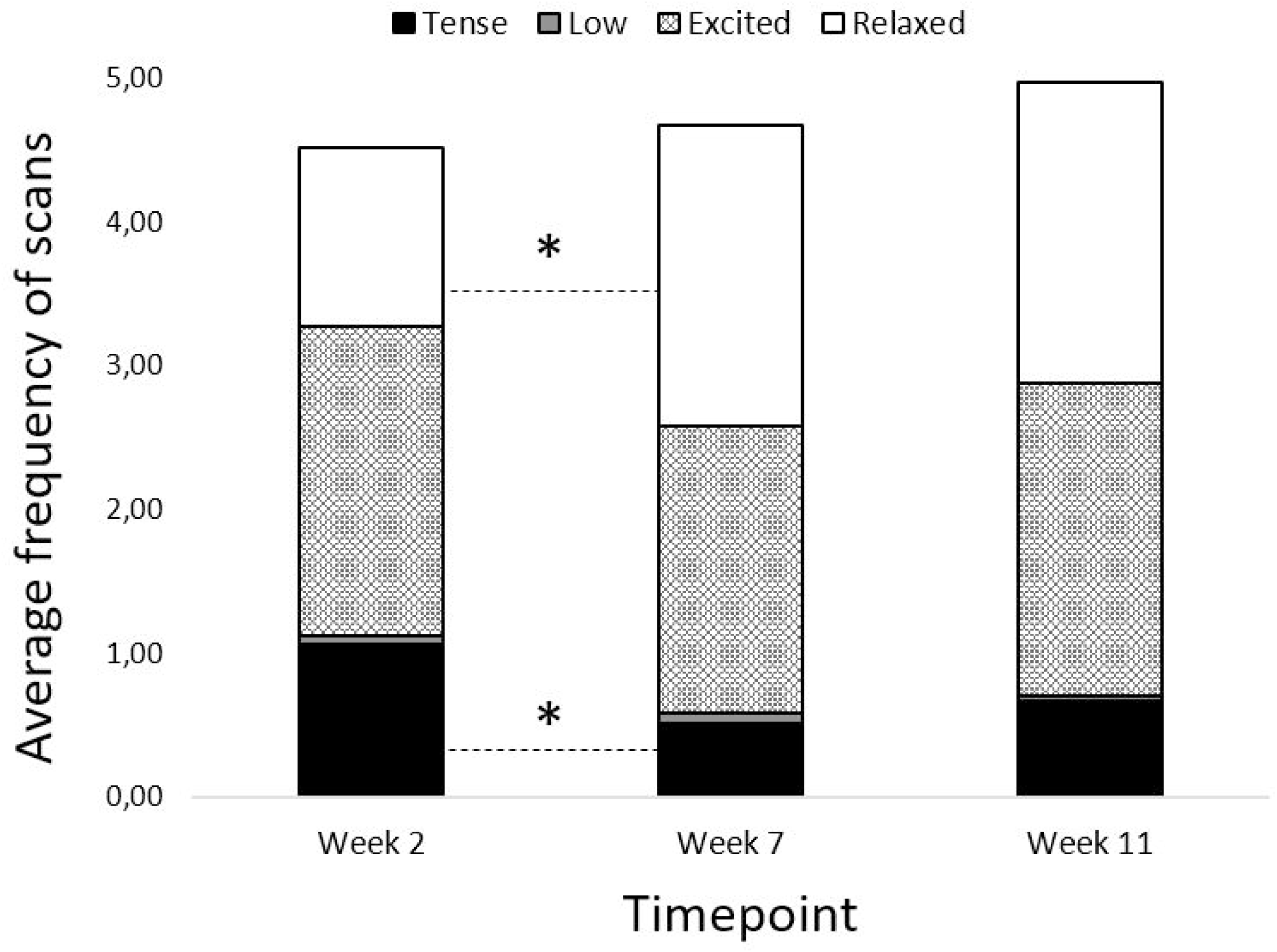
Average frequency of scans in the different behavioral states during the training sessions at the prisons in Week 2, Week 7 and Week 11 (N=33). The cases where the bars do not reach the maximum (5) correspond to classification of the behavioral state as ‘Unknown’ (see Appendix 6). *stands for statistically significant differences.

**Figure 6.**
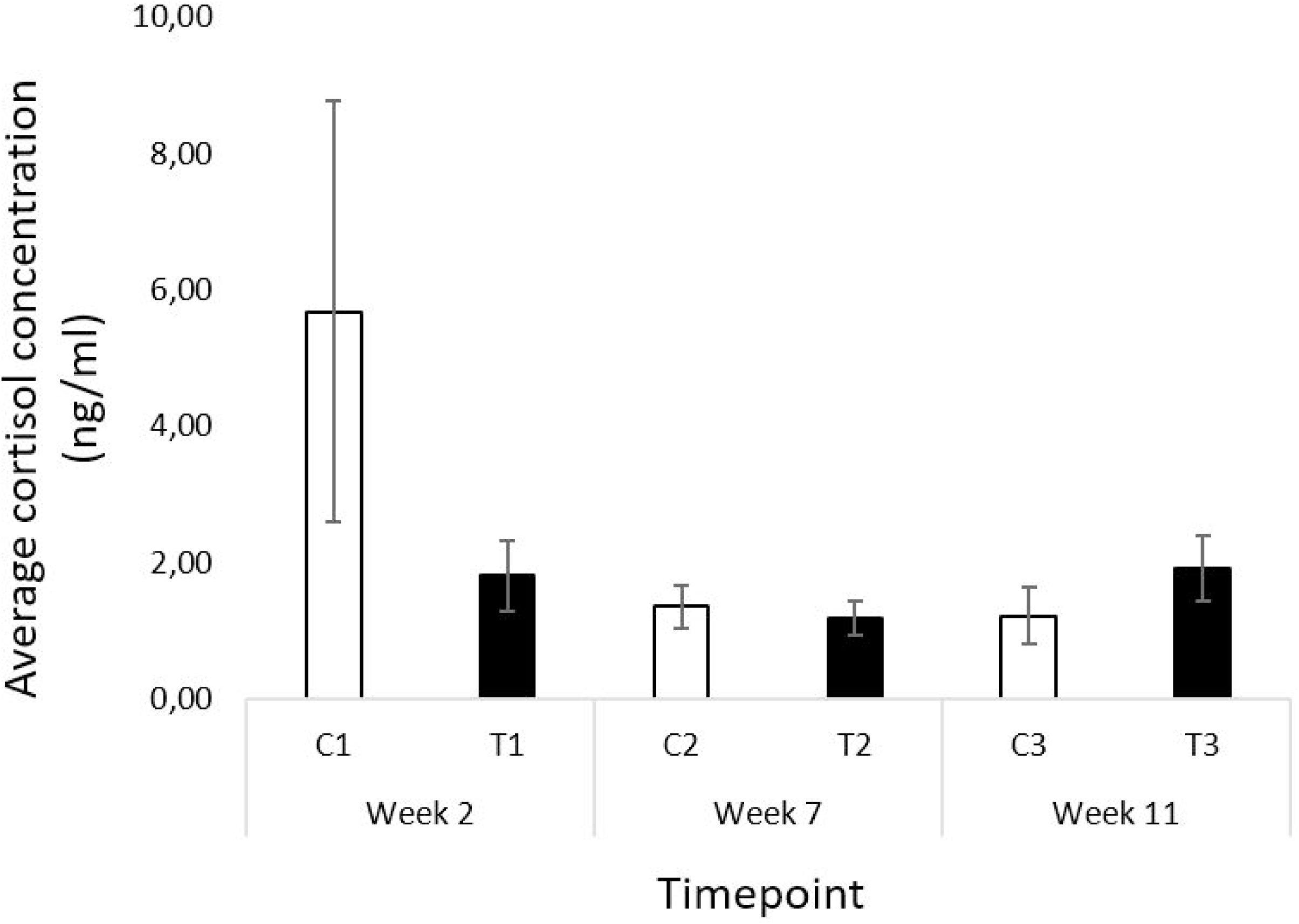
Average salivary cortisol concentration (ng/ml) on training days at the prisons (T) and on non-training days at the shelter (C), across three time-points (Week 2, Week 7 and Week 11, N=34).

#### 3.2.3. Salivary Cortisol

No differences were found between pre-program baseline cortisol levels (B) and control cortisol levels obtained on non-training days at the shelter in the course of the PBDTP (C1 to C3) [B vs C1: t(10)=1.85, p=0.094; B vs C2: t(11)=-0.35, p=0.732; B vs C3: t(10)=2.10, p=0.062]. Following these results, we concluded that C levels could be used as appropriate controls against which to compare the cortisol levels at the prisons in training days (T). Hence, C levels were included in the main cortisol model as control values. Thereafter, we found no effects of control cortisol level C [F_(1,33)_=0.20, p=0.660], time-point [F_(2,33)_=1.41, p=0.258] or prison [F_(2,33)_=0.00, p=0.996] in cortisol level T. Figure 3 shows the average cortisol concentrations for C1-C3 and T1-T3. Moreover, we found no differences between pre-program baseline cortisol levels (B) and post-program cortisol levels (F) [t(9)=1.33, p=0.217].

#### 3.2.4. Cognitive Bias

The (adjusted) latency to reach the food bowl in the ambiguous locations did not change significantly from pre- to post-program [Pre-program: 46.56±14.96 vs post-program: 47.03±22.70; F_(1,15)_=0.08, p=0.782].

### 4.1. Discussion

This paper presents a comprehensive study of how dogs are affected by participating in a PBDTP, combining a battery of socialization and handling and basic training indicators with behavioral, physiological and cognitive indicators of welfare. Our results show an overall improvement in the basic training and socialization and handling skills of the dogs as a consequence of participating in the PBDTP, whereas there was no indication of lasting stress.

Participation in the 12-week training program resulted in an improvement in sociability towards humans, playful behavior, and in almost all education categories that were tested. A significant improvement was seen even for BET categories where the average performance was around 90% already at the start of the program, i.e. where the dogs already displayed good basic training skills prior to participating in the PBDTP.

Changing handlers has been found to reduce performance [33], and inexperienced handlers are often not timing reinforcement accurately [34], which can impact learning negatively. However, our study found that the training was effective even though the dogs were trained by several handlers who had no previous dog training experience and hence were initially unskilled. The dogs participating in this program were from a shelter and were available for adoption after completion of the program. Given that behavioral issues are the most frequently cited reason for relinquishment or rehoming [35], there is reason to believe that the improvement in socialization and handling and basic training may help the dogs be successfully adopted i.e., integrate well and not be returned to the shelter or cause extreme stress to their adopters.

Among all tested socialization and handling and education categories, dogs did not improve only on their scores for Dog Sociability (sociability towards other dogs) and Recall (coming when called). Socializing dogs with conspecifics was not part of the designed training program and, despite the fact that training sessions could have contributed to improve dog sociability through desensitization and counterconditioning (sessions were held in groups of 6 dogs, in a way that dogs were exposed to the presence of others from a distance that was comfortable to them, and they were receiving food and other potentially appetitive stimuli such as positive human interactions), it may be the case that targeted training is needed for this specific behavior to improve [36]. Regarding Recall, this behavior was trained during the program for distances up to the 2 meters as allowed by the leashes on which the dogs were kept during the training sessions. The 10-meters distance dogs had to cover during testing was never directly trained, and distance is advised by dog training professionals as an important variable to work on when training a dog [37].

The second important factor here is the welfare of the participating dogs. Participating in the program subjected the dogs to several unfamiliar and potentially stressful experiences. Transport, at least if long (≥50 minutes) has been shown to affect dog welfare negatively [38]. The participating dogs were transported by car to and from the prisons twice a week, at a driving distance of approximately 23 to 35km, and they were (to our knowledge) initially not used to neither crating nor transport. The training itself was exclusively reward- based, thus avoiding methods known to cause stress [32], but the fact that they took place in an initially unfamiliar place is a potential source of stress, as is the handling by several different and initially completely inexperienced handlers, and the proximity to other dogs. Despite these potentially stressful factors, there was no evidence that the dogs were overall negatively affected. Participating in the training program did not affect levels of salivary cortisol, nor performance in the cognitive bias test of affective state, and behaviors indicative of stress were generally rare during training sessions.

At the first time-point (week 2 of training in the prisons), lip licks were more frequent and dogs were often seen in a tense state and less often in a relaxed state compared to later time-points. The reduction in stress-related behaviors between the first (week 2 of training) and the second (week 7 of training) time-points shows that the dogs did not take long to habituate, and that the combination of transport and training did not result in long-lasting stress.

As observed by Cobb et al. [39], there is a large amount of intra-individual and inter- individual variability and external variables that could influence salivary cortisol concentration. In line with this, in the present study, two dogs had much higher C1 cortisol values than the remaining (11) animals, which resulted in the high variability observed for this time-point. The reasons underlying the elevated values for these animals are beyond our knowledge, as no demographic factor including sex, age, weight, neuter status or time spent in the kennel before study commencement differentiates these animals from the remaining sample. We have tried to control for these variations in our experimental design by comparing cortisol concentration only intra-individual, and by standardizing collection timing and conditions as much as possible. However, the sample size for this welfare measure turned out to be much smaller than initially planned. Due to several samples not having enough saliva volume for laboratory analysis and because we used a repeated-measures design, we ended up with only 7 to 14 subjects per comparison, despite having collected saliva samples of 34 animals. This small sample size thus prevents any strong conclusion from being drawn. However, the analysis of stress-related behaviors during training sessions, for which the sample size was only one less than the intended, suggest that participating in the PBDTP was not stressful for the animals. In fact, of the six behaviors analyzed, five were extremely rare (absent in 64-96% of the observations), and these same behaviors have been shown to occur frequently in a stressful training context [32].

The present findings are in line with Leonardi et al [25] and Hennessy et al [24], in that all the three studies found improvements in dog training performance after participating in a PBDTP, and in line with Hennessy et al [24] in that both studies found no changes in cortisol levels from before to after the program. Leonardi et al [25] reported improvements in dog welfare, with dogs showing more relaxed behaviors in their kennels after the PBDTP. In the present study, we also found dogs to become more relaxed and less tense across the program. However, our assessments were performed during training sessions in the prisons and not in the shelter as in Leonardi et al [25], and hence are not directly comparable. Moreover, in Leonardi et al [25], shelter staff rated dogs as more playful, friendly and relaxed, and less excitable, frustrated, noisy, reactive or vigilant after participating in the PBDTP. Despite further suggesting that dog welfare improved with the PBDTP, these results need to be interpreted with caution because they rely on subjective ratings by shelter staff who were not blind to the treatment (i.e., participation in the PBDTP). Importantly, the present study only used objective and quantitative welfare indicators and evaluated welfare both within and outside the training context, thus providing a more robust assessment of the welfare of dogs participating in PBDTPs. Taken together, data available up to date suggest that participation in PBDTPs has, at least, no negative impact on dog welfare.

The present study provides comprehensive data on dog behavior and welfare as a result of participating in a PBDTP. The fact that it included a sizeable sample (42 dogs) and three different prison establishments with different sets of inmates and supervising dog trainers are further strengths of the study. In the vast majority of PBDTPs, the dogs live full- time at the prisons, as opposed to living in shelters or other facilities and being transported to the prisons for the training sessions as in the case of Pelos2. Besides animal housing, PBDTPs vary in the number of inmates with whom each dog is paired, number of participants, criteria for participation, length of participation, or the amount of time per day that the participants spend with the dogs [16]. It is still unclear how different aspects of PBDTPs may affect their efficacy and impact on animal welfare, and hence the generalization of the present results is somewhat limited. Behavior during training sessions could only be analyzed through direct observations, as video recordings would have raised issues with protecting the identity of a vulnerable population of human participants. To minimize the risk of bias and maximize the reliability of the direct observations, we made an effort to train and calibrate the two observers before they started to collect data. As regards the tests for dog socialization and handling and basic training, the exercises in the Basic Education Test were originally designed to be performed by the dog’s owner or regular handler, and the fact the dogs here were tested with research personnel may have affected their performance.

The improvement in basic training and socialization and handling skills, and the absence of indicators that the participation was stressful for the dogs suggest that the program can be deemed a success for the long-term benefits it is likely to bring to the dogs and their future adopters. In future research, it would be important to evaluate to what extent the improvement in dog behavior actually translates into an increase in successful adoptions.

## Author Contributions

Parizad Baria-Unwalla: Conceptualization, Data curation, Formal analysis, Investigation, Methodology, Validation, Visualization, Writing – original draft, Writing – review & editing; Gabriela Munhoz Morello: Formal analysis, Writing – review & editing; Maria Queiroz: Conceptualization, Data curation, Methodology; I. Anna S. Olsson: Conceptualization, Funding acquisition, Project administration, Resources, Supervision, Validation, Writing – original draft; Ana Catarina Vieira de Castro: Conceptualization, Data curation, Formal analysis, Investigation, Methodology, Project administration, Supervision, Validation, Visualization, Writing – original draft.

## Acknowledgements

We would like to thank the entire DTC team for making this study possible, especially Luca Conde and Inês Vieira, for being so enthusiastic about this collaboration since the beginning. We are also deeply grateful to the DTC trainers involved in the Pelos2 program, particularly Rita Pinto, Marco Saraiva, André Carneiro and José Xavier, as well as to the DTC psychologist Silvia Sousa, who facilitated the collection of live behavior data at the prisons, ensuring our research did not interfere with their training schedules.

Our gratitude goes to the staff at the prison establishments of Paços de Ferreira, Vale do Sousa, and Santa Cruz do Bispo, as well as the shelter staff at PATA, for their cooperation and flexibility, which greatly contributed to the successful completion of this study.

We extend our heartfelt thanks to Sofia Morais and Patricia Fachada for their invaluable assistance with data collection at the shelter. We would also like to express our appreciation to the students from the Abel Salazar Biomedical Sciences Institute at the University of Porto and the University of Edinburgh, whose voluntary efforts in scoring videos strengthened the robustness of our results.

Finally, we are particularly grateful to the editors and the two anonymous reviewers at Animal Behavior and Cognition, whose constructive feedback greatly improved our study protocol. We also thank Ana Maria Valentim for her valuable input on our final manuscript.

