## Appendices for "What’s in it for the dogs? Assessing the outcomes of a prison-based dog training program from an animal behavior and welfare perspective"

### Appendix 1 – Demographic data for the participating dogs

| Batch | Dog Name | Sex | Age (Y) | Weight (kg) | Sterilized | PATA Entry |
|---|---|---|---|---|---|---|
| 1 | Lily | F | 3 | 20-25 | Yes | 20/07/2021 |
| 1 | Júlio | M | 7 | 20-25 | Yes | 04/11/2020 |
| 1 | Totti | M | 5 | 10-15 | Yes | 17/11/2021 |
| 2 | CucaW | F | 4 | 30-35 | Yes | 13/09/2022 |
| 2 | CucaB | F | 3 | 25-30 | Yes | 02/11/2021 |
| 2 | Mila | F | 5 | 20-25 | Yes | 19/11/2019 |
| 2 | Bongo | M | 2 | 20-25 | No | 08/09/2022 |
| 2 | Farofa | M | 1.5 | 15-20 | Yes | 01/04/2022 |
| 2 | Ralph | M | 3 | 25-30 | Yes | 18/07/2022 |
| 3 | Nino | M | 3 | 25-30 | Yes | 06/12/2022 |
| 3 | Clooney | M | 6 | 25-30 | Yes | 05/12/2022 |
| 3 | Oliveirinha | F | 3 | 15-20 | Yes | 19/09/2022 |
| 3 | Terra | F | 8 | 20-25 | Yes | 20/08/2021 |
| 3 | Bóris | M | 8 | 25-30 | Yes | 14/10/2021 |
| 3 | Belinha | F | 8 | 15-20 | Yes | 31/01/2017 |
| 3 | Noa | F | 3 | 25-30 | Yes | 21/12/2022 |
| 3 | Max | F | 4 | 15-20 | No | 25/07/2022 |
| 3 | Mel | F | 2 | 15-20 | Yes | 27/09/2022 |
| 3 | Jonas | M | 6 | 15-20 | Yes | 20/10/2022 |
| 3 | Loki | M | 2 | 20-25 | Yes | 30/10/2022 |
| 3 | Tomás* | M | 1 | 10-15 | Yes | 21/10/2022 |
| 3 | Alex* | M | 1 | 10-15 | Yes | 21/10/2022 |
| 3 | Sequeira | M | 5 | 30-35 | Yes | 21/11/2022 |
| 3 | Luck** | M | 1 | 20-25 | No | 13/10/2022 |
| 3 | Caviar | M | 1 | 20-25 | No | 16/08/2022 |
| 3 | Santa | F | 8 | 10-15 | Yes | 25/10/2021 |
| 4 | Poupas** | M | 2 | 5-10 | Yes | 10/03/2023 |
| 4 | Soneca | M | 2 | 20-25 | No | 28/10/2022 |
| 4 | Bully | M | 8 | 30-35 | Yes | 22/11/2021 |
| 4 | Panda | M | 2 | 20-25 | No | 28/10/2022 |
| 4 | Tiger | M | 2 | 20-25 | Yes | 28/10/2022 |
| 4 | Lázaro | M | 8 | 20-25 | Yes | 28/10/2022 |
| 4 | Ottis** | M | 3 | 20-25 | Yes | 20/02/2023 |
| 4 | Afonso | M | 3 | 5-10 | Yes | 22/03/2023 |
| 4 | Fox | M | 8 | 15-20 | Yes | 23/11/2015 |

| | | | | | | |
|---|---|---|---|---|---|---|
| 4 | Rex | M | 5 | 40-45 | Yes | 21/03/2023 |
| 4 | Platt | M | 8 | 15-20 | Yes | 09/07/2018 |
| 4 | Surya | F | 8 | 20-25 | Yes | 06/01/2023 |
| 4 | Jack | M | 2 | 20-25 | Yes | 22/04/2023 |
| 4 | Ramada | M | 1 | 15-20 | No | 17/03/2023 |
| 4 | Márcio** | M | 3 | 20-25 | Yes | 31/03/2023 |
| 4 | Tito | M | 8 | 25-30 | Yes | 05/01/2021 |

*Unmanageable on leash during pre-program BET; not tested with BET at this stage

**Did not allow saliva collection; excluded from cortisol sampling

**Appendix 2 – PBDTP Training Plan Summary (developed and implemented by DTC Social)**

Each inmate participated in 24 sessions (2 sessions per week), each of 45-minute duration, broadly outlined below.

**Session 1** was an introduction to DTC Social, their work philosophy, the Pelos2 project, and the role the inmates were expected to fulfil. In this session, the inmates interacted with DTC dogs (not the dogs from the shelter who are the main participants in the program). They learned about dog breeds, were taught how dogs like to be petted and got to pet the animals.

**Session 2** was for inmates to understand the differences between dogs, breed specific characteristics, understand canine wellbeing and their needs, and reflect on how the program could help them and the dogs. In this session, they got to work with some equipment like leashes, snack packs, food rewards, brushes, etc. They were still working with DTC dogs.

**Session 3** was the first session where the inmates were introduced to the shelter dogs they would be training (i.e. the participants of this study). Each inmate was paired with a dog and given rules for interacting properly with them. They worked primarily on attachment activities like walking the dogs, feeding them, and petting them. They rewarded their dog with food when the dog responded to their name (recall). The dogs were on leash at all times.

**Session 4** was focused on inmates understanding the basic concepts of training and communication, both verbal and non-verbal, and basic concepts of animal welfare. They learned about different leashes and harnesses, and how to safely put them on. They also learned how to lure the dogs and reward them for responding appropriately.

**Session 5** aimed to teach inmates to understand the dogs' body signals, develop a general understanding of calming signals, and apply this understanding to assess dog welfare. They also learned how to help the dogs overcome "obstacles" (doors, stepping over a bar on the floor, or overcoming a specific fear) using luring and positive reinforcement.

**Session 6** aimed to teach inmates to understand the signs and origins of fear in dogs, to learn about stimulus and response in behavior, concepts of environmental management, importance of leash walking without pulling, and how to identify and address compromised wellbeing in dogs.

**Session 7** was to understand the difficulties that may arise in the communication process, and the concept of assertive communication. They continued to work on leash walking exercises and practice luring with food rewards to guide the dogs to move from side to side, turn around, go over items on the floor, and respond to the recall.

**Session 8** focused on inmates understanding the concepts of marker (e.g., a verbal "good boy"), the importance of reinforcement for learning, and the importance of timing. Inmates taught their dogs to sit (guiding through luring) with a palm-up hand gesture. They learned the process of teaching a behavior and how to help the dogs when they present learning difficulties.

**Session 9** aimed to understand the importance of play, explore the different ways of play, and recognise dogs' own play preferences. Inmates performed exercises for improving posture and clarity in (cue) gestures as demonstrated by the trainers. They tried different types of play with their dogs including tug, fetch or ball toy.

**Session 10** was to learn how to recognize and overcome communication difficulties, how to focus on two things at the same time (dog and real-life situations) through role-play with the trainers, and how to teach a dog to lay down through luring with a standard hand gesture (hand outstretched, palm facing down, downward movement).

**Session 11** was a mid-term evaluation where inmates demonstrated their training skills with the dog, based on the Canine Good Citizen – Basic Education Test (AKC, 2023). They received feedback and recommendations from trainers on aspects that needed improvement.

**Session 12** focused on inmates understanding the basics of dog body signs and communication. They learned about stereotypies and different body postures. The dogs were walked on leash in pairs, crossing each other, and rewarded for remaining calm. The importance of "doing nothing" was emphasized and the inmates had some free time to use as they wish.

**Session 13** focused on training different skills simulating real-life situations, and the importance of waiting skills. Skills like sit, lie down, recall, or staying still were practiced in imaginary situations like pedestrian crossings at traffic lights. Inmates reflected and discussed situations that were more difficult for their dogs to remain calm/waiting.

**Session 14** was for inmates to recognize compromised wellbeing in dogs and to know how to act in such situations. They learned about relaxation activities to deal with stress and anxiety, different kinds of situations that can cause dogs to fear, and the importance of nose work as a natural and relaxing behavior. They also practiced training in the presence of distractions like noise and strangers.

**Session 15** focused on recognizing the importance of timing and delivery of reinforcement, understanding motivation in dogs and using external motivators, and concentrating on multiple tasks simultaneously. Inmates trained the dogs to stay still, using positive reinforcement, in increasing level of difficulty for the animals.

**Session 16** emphasized the importance of relaxation in building a balanced dog. The exercises previously trained were repeated in a calm manner. Nose work games were practiced. Inmates created relaxing games for their dogs, and learned to be together with caresses, brushing, or even doing nothing, as per the dogs' preference.

**Session 17** explained the differences that come with changing a handler, including advantages and challenges. The inmates worked with a different dog (not the one they were normally paired with) and worked on all previously trained exercises using the same gestures. Inmates exchanged observations and notes with each other.

**Session 18** emphasized the importance of habituation in the adaptation of the dog. The inmates trained the dogs to accept a muzzle, in small steps. Inmates also got time to choose whether to train, play, or relax for a few minutes at the end of the session.

**Session 19** was dedicated to preparation for the final assessment. Inmates were told about the evaluation criteria of the test applied to the dogs. They reviewed all exercises and practiced a Canine Good Citizen (AKC, 2023) test in a mock setup. The group reflected on the advantages of the program for future adopters, and their own contribution to the evolution of the dogs.

**Session 20** revised the importance of the habituation process, and extended the concepts used for training muzzle acceptance to other everyday situations like walking past dogs, people, etc. while remaining calm. A variation of the "musical chairs" game was played by the inmates with their dogs.

**Session 21** revised the concept of motivation in dogs. Inmates practiced all the exercises learnt using different reinforcement (food, different toys) to explore their dogs' preferences. The exercises were then repeated using social reinforcement (petting and/or praising) only.

**Session 22** focused on preparation for the final assessment. All the concepts from the previous sessions were reviewed and the Canine Good Citizen (AKC, 2023) obedience competition was simulated. Trainers helped inmates solve difficulties with their individual dogs.

**Session 23** aimed at reflecting on the journey of the dogs and the inmates. They came up with ideas for new behaviors to teach their dogs (like pawing, rolling over, etc.), learn how to teach them those skills, and practice new behaviors with the guidance of the trainers.

**Session 24** marked the end of the program. Inmates were explained about what life will be for the dogs after the program, outreach platforms, adoption, etc. The session was a personal time for inmates to say goodbye to their dogs in the way they choose. The session ended with group reflection, presentation of memories, and a message of motivation for the future.

**Appendix 3 – Basic Education Test (BET) Protocol and Scoring System**

The test protocol is an adaptation from the Canine Good Citizen Test[1] and was piloted at PATA in a previous study[2]. The test was conducted in an indoor room at the shelter measuring approximately 12m x 8m. Three experimenters participated in the tests: the 'handler', who handled and performed the exercises with the dogs (MQ), the 'evaluator', who scored the dogs' behavior live (ACVC) and a third person who was responsible for video recording the tests with a video camera on a tripod (PB). The dogs were tested in the same order in pre-program and post-program assessments.

| **Scoring Description** | **Score** |
|---|---|
| **Item 1 – Accepting a Friendly Stranger**: The dog is on leash with the handler and the evaluator approaches them, stopping at a distance of 1 meter. The evaluator greets the handler in a friendly manner, ignoring the dog (for example, "Hello, how are you? Nice to see you here!"). In this exercise, the evaluator does not interact with the dog. The handler must not tighten or retract the leash to prevent the dog from moving forward/jumping or moving away/retracting during the exercise. | |
| Dog is friendly and calm and does not jump or lunge forward to greet the evaluator. | 3 |
| Dog is friendly but excited and tries to jump or lunge forward to greet the evaluator. | 2 |
| Dog retreats from the evaluator, showing fear. | 1 |
| Dog shows signs of aggression towards the evaluator. | 0 |
| **Item 2 – Waiting Politely for Petting**: To begin the exercise, the evaluator asks, "May I pet your dog?". Next, the evaluator approaches the dog and tries to pet it on the head and body. The handler may talk to the dog throughout the exercise. After petting the dog, the evaluator backs away. The handler must not tighten or retract the leash to prevent the dog from moving forward/jumping or moving away/retracting during the exercise. | |
| Dog is friendly and calm and does not jump or lunge forward towards the evaluator. | 3 |
| Dog is friendly but excited and tries to jump or lunge forward towards the evaluator. | 2 |
| Dog retreats from the evaluator, showing fear. | 1 |
| Dog shows signs of aggression towards the evaluator. | 0 |
| **Item 3 – Walking on a Loose Leash**: There will be a pre-planned course of 30 meters, with a right turn, a left turn and a 180º turn. The handler may talk to the dog during the course. It is not required that the dog walks besides the handler, and the dog is allowed to sniff as long as it does not pull the leash and performs the changes of direction appropriately. The handler must not tighten or retract the leash during the exercise. | |
| Dog walks all the way without pulling. | 3 |
| Dog pulls occasionally, up to three times. | 2 |
| Dog pulls often, more than three times. | 1 |
| Dog pulls all the way. | 0 |

[1]AKC American Kennel Club. Canine Good Citizen Test. [Online]. [cited 2024 October 16]. Available from: https://www.akc.org/products-services/training-programs/canine-good-citizen/.

[2]Vieira de Castro AC, Malone M, Casaca M, Blenkuš U, Gomes L, Baria-Unwalla P, Olsson IAS. Evaluating basic obedience and temperament tests for shelter dogs participating in a training program. Journal of Veterinary Behavior. Under review.

| | |
|---|---|
| **<u>Item 4 – Walking Through a Crowd</u>**: The dog and handler walk around and pass close to three/four people (some are still, some are moving, in order to simulate a public place) during 30 seconds. The evaluator can be counted as one of the people in the crowd. The dog may show some interest in the people, but it should continue to walk with the handler. The dog may even sniff people briefly but must, however, move on right away. | |
| Dog is friendly, calm, walking on a loose leash and does not jump or lunge forward towards the strangers. | 3 |
| Dog is friendly but excited and tries to jump or lunge forward towards the strangers. | 2 |
| Dog retreats from the strangers, showing fear. | 1 |
| Dog shows signs of aggression towards the strangers. | 0 |
| **<u>Item 5 – Sit, Down, and Staying in Place</u>**: Before the exercise begins, the dog's leash is replaced by a 10-meter leash. For the stay in place exercise, the handler can choose to leave the dog sitting or lying down. The handler cannot use force or intimidation to place the dog in either position. When instructed by the evaluator, the handler cue the dog to sit, then to lie down and, finally, to stay (in a sitting or lying position, at the handler's choice), and walks away 10 meters in a straight line. When the distance of 10 meters is reached, the handler immediately returns to the dog. The whole process must be done at a normal pace. The cues can be either verbal or gestural. | |
| Dog performs the three exercises at first attempt (even if the dog changes position in the third exercise, as long as it remains in place). | 4 |
| Dog performs the three exercises with two attempts each. | 3 |
| Dog fails to perform one of the exercises (two attempts per exercise are allowed). | 2 |
| Dog fails to perform two exercises (two attempts per exercise are allowed). | 1 |
| **<u>Item 6 – Coming When Called</u>**: When starting the exercise, the dog is left free to explore the environment (loose or with a 10-m leash, depending on the safety of the test area). When instructed by the evaluator, the handler calls the dog. The exercise is considered finished when the dog approaches the handler and the latter grabs the collar and/or places the leash on. The handler can use body language and encouragement when calling the dog. | |
| Dog fails to perform the three exercises (two attempts per exercise are allowed). | 0 |
| Dog succeeds to come when called at first attempt. | 2 |
| Dog succeeds to come when called at second attempt. | 1 |
| Dog fails to come when called within two attempts. | 0 |
| **<u>Item 7 – Supervised Separation</u>**: When instructed by the evaluator, the handler passes him the dog's leash and moves away, remaining out of the dog's field of vision for 3 minutes. If the dog begins to show many signs of anxiety or discomfort, the exercise should be terminated. | |
| Dog stays calm and under control. | 2 |
| Dog walks back and forth and looks for the handler but does not show signs of distress. | 1 |
| Dog shows signs of distress (barking, howling, panting vigorously, pacing, and pulling). | 0 |

**Appendix 4 – Temperament Test (TT) Protocol and Scoring System**

The test protocol was based on the Temperament Test[3,4] and was piloted at PATA in a previous study[2].

Subtests 1-8 were conducted in the individual dogs' home enclosures at the shelter, after separating them temporarily from other dogs they might be housed with. Subtests 10-18 were conducted in the same indoor room where the BETs were conducted. Subtests 9 and 19 were recorded between the home enclosures and the indoor room, and included walking through other indoor areas in the building and outdoor areas where the dogs were housed.

Three experimenters participated in the tests: the 'handler', who handled the dogs throughout the test (MQ for pre-program assessment and ACVC for post-program assessment), the 'evaluator', who scored the dogs' behavior live (ACVC for pre-program assessment and MQ for post-program assessment) and a third person who was responsible for video recording the tests with a video camera on a tripod (PB). The dogs were tested in the same order in pre-program and post-program assessments.

| Test Component | Subtest | Experimenter's Behavior | Scoring Description | Score |
|---|---|---|---|---|
| Behavior in Kennel | 1 – Observation from a distance | Out of the dog's sight, records the dog's position in kennel (30s)<br><br>(Consider the position where the dog remains the longest during the observation period) | Front | 3 |
| | | | Cannot determine whether front or centre | 2.5 |
| | | | Centre | 2 |
| | | | Cannot determine whether centre or back | 1.5 |
| | | | Back | 1 |
| | | | Out of sight/Impossible to determine where the dog remains the longest | 0 |
| | 2 – Stereotypical behavior | Records the presence of stereotypies (pacing, circling, jumping, etc.) (30s) | No | 1 |
| | | | Yes | 0 |
| Human Sociability | 3 – Kennel approach | Approaches kennel in a neutral posture, no eye contact, and stops in front of the fence facing sideways (30s)<br><br>Body posture:<br>• friendly (tail wagging, non-aggressive barking, calm, etc.)<br>• neutral (dog is still, neither threatening nor friendly behavior) | Friendly, approaches the experimenter seeking contact | 9 |
| | | | Friendly but not asking for contact | 8 |
| | | | Approaches with excitement, jumps/struggle to calm down | 7 |
| | | | Neutral, calm approach | 6 |
| | | | Neutral and still, looks at the experimenter | 5 |
| | | | Neutral but avoiding contact | 4 |
| | | | Fearful, approaches in a low posture | 3 |
| | | | Fearful and still | 2 |

[3] Valsecchi P, Barnard S, Stefanini C, Normando S. Temperament test for re-homed dogs validated through direct behavioral observation in shelter and home environment. Journal of Veterinary Behavior. 2011; 6(3): 161-177.

[4] Barnard S, Kennedy D, Watson R, Valsecchi P, Arnott G. Revisiting a previously validated temperament test in shelter dogs, including an examination of the use of fake model dogs to assess conspecific sociability. Animals Special Issue: The Welfare of Cats and Dogs. 2019; 9(10): 835.

| | | | | |
|---|---|---|---|---|
| | | • fearful (crouched posture, low tail, shaking, whimpering, etc.)<br>• aggressive/threatening (barking, growling, lunging towards experimenter, etc.) | Fearful, showing signs of avoidance (withdraws, tries to hide) | 1 |
| | | | Aggressive/threatening | 0 |
| | 4 – Side crouch | Crouches down, side on, near the fence, talks calmly to the dog (30s) | Friendly and confident | 3 |
| | | | Neutral, less confident | 2 |
| | | | Fearful | 1 |
| | | | Aggressive/threatening | 0 |
| | 5 – Stroking through the fence | Calls the dog, talks gently, attempts to stroke dog through fence (30s) | Friendly and confident | 3 |
| | | | Neutral, less confident | 2 |
| | | | Fearful | 1 |
| | | | Aggressive/threatening | 0 |
| | 6 – Entering kennel | Walks into kennel and closes the door. Stands still, arms at side, ignoring the dog (30s) | Approaches the experimenter | 2 |
| | | | Does not move | 1 |
| | | | Moves away | 0 |
| | 7 – Physical contact | Bending slightly, calls the dog in a relaxed manner, waiting for the dog to make contact (30s) | Friendly, approaches the experimenter seeking contact | 9 |
| | | | Friendly but not asking for contact | 8 |
| | | | Approaches with excitement, jumps/struggle to calm down | 7 |
| | | | Neutral, calm approach | 6 |
| | | | Neutral and still, looks at the experimenter | 5 |
| | | | Neutral but avoiding contact | 4 |
| | | | Fearful, approaches in a low posture | 3 |
| | | | Fearful and still | 2 |
| | | | Fearful, showing signs of avoidance (withdraws, tries to hide) | 1 |
| | | | Aggressive/threatening | 0 |
| Behavior on Leash | 8 – Placing on leash | Standing inside the kennel, shows the leash to the dog (30s). Then attempts to put it on. | Dog is calm and confident, and leash is easily put on | 2 |
| | | | Dog is fearful, but leash is easily put on | 1.5 |
| | | | Dog is excited or reluctant, placement of leash is relatively difficult | 1 |
| | | | Dog is too excited or too reluctant, placement of leash is very difficult | 0.5 |
| | | | Dog reacts aggressively or with extreme fear and it is impossible to put the leash on | 0 |
| | | | Easy to handle | 2 |

| | | | | |
|---|---|---|---|---|
| | 9 – Walking on leash | Walks dog from kennel toward testing area (30 – 60s) | Excited, pulls, but is manageable | 1.5 |
| | | | Scared, reluctant to walk | 1 |
| | | | Pulls a lot or refuses to walk, very difficult to handle | 0.5 |
| | | | Pulls a lot or refuses to walk, impossible to handle | 0 |
| | | A 10-m leash is placed, leaving it to drag on the ground. The dog is allowed to explore the room for 2 min. Then proceed with the following steps: | | |
| Human Sociability | 10 - Handling | 1. General handling - strokes dog from head to tail (with a fake hand if needed), and tries to move and lift one of the front paws (15s) | Calm/confident | 2 |
| | | | Uncomfortable (freezes, licks lips, may show a low posture), but allows handling | 1 |
| | | | Aggressive or extremely fearful, does not allow handling/growls | 0 |
| | | 2. Brushing – head and body (15s) | Calm/confident | 2 |
| | | | Uncomfortable (freezes, licks lips, may show a low posture), but allows brushing | 1 |
| | | | Aggressive or extremely fearful, does not allow brushing/growls | 0 |
| | | 3. Put muzzle on (15s) | Calm/confident | 2 |
| | | | Uncomfortable (freezes, licks lips, may show a low posture), but allows muzzle placement | 1 |
| | | | Aggressive or extremely fearful, does not allow muzzle placement | 0 |
| Cognitive Skills | 11 - Attention | Standing in front of the dog, shows a treat with the hand. Then closes the hand and brings it to chest. The dog must follow the hand movement and keep looking (3s). | Holds gaze for 3 sec | 1 |
| | | | Jumps at hand or looks away | 0 |
| | 12 – Problem solving | Shows the dog a treat and evaluates its interest in the food. Showing a second treat and making sure the dog is watching, places the treat in a bowl facing down. Allows the dog to try to retrieve the treat (30s). | Solves the task | 3 |
| | | | Attempts to solve the task | 2 |
| | | | Interested but does not attempt to solve the task | 1 |
| | | | Not interested in food or task | 0 |
| Playfulness | 13 – Squeaky toy | Show the toy to the dog, play the squeak and throw it close to him | Plays with the toy | 5 |
| | | | Displays playful behavior not directed at the toy | 4 |
| | | | Approaches confidently | 3 |

| | | | | |
|---|---|---|---|---|
| | | | Approaches with caution | 2 |
| | | | Shows no interest | 1 |
| | | | Dog is scared | 0 |
| | 14 – Ball toy | a. Show the dog the ball, allowing it to sniff and throws the ball away | Plays with the ball | 1 |
| | | | Does not play with the ball | 0 |
| | | b. If the dog brings the ball back, throw it once again | Was the ball brought back?[a] | |
| | | | Yes | 2 |
| | | | No | 1 |
| | | | Reluctant to let the ball go/growls | 0 |
| Reactivity | 15 – Food bowl removal | With dog on leash, provides a full bowl of dry food. Dog approaches food, few mouthfuls allowed. Then uses fake hand to pull bowl away. | Allows the bowl to be removed without showing signs of tension, discomfort, or fear, ignores the experimenter. | 3 |
| | | | Dog looks anxious and eats faster or moves away | 2 |
| | | | Shows no interest in food | 1 |
| | | | The dog guards the bowl, growling. It is impossible to remove the bowl. | 0 |
| Dog Sociability | 16 – Fake dog approach | a. Small fake dog (similar to a Jack Russel Terrier – Melissa & Doug brand): With the dog on a leash, approach the fake dog | Calm, confident and friendly | 2 |
| | | | Shows no interest, becomes overly excited or fearful/reluctant | 1 |
| | | | Tense/growls | 0 |
| | | b. Big fake dog (Similar to a Labrador Retriever – Melissa & Doug brand): With the dog on a leash, approach the fake dog | Calm, confident and friendly | 2 |
| | | | Shows no interest, becomes overly excited or fearful/reluctant | 1 |
| | | | Tense/growls | 0 |
| Reactivity | 17 – Reactivity to auditory stimulus | With the dog on a leash, approximately 3 m from the dog, without establishing visual contact, the experimenter plays a bicycle horn | Curious, wants to explore stimulus | 3 |
| | | | Does not show any reaction | 2 |
| | | | Fearful, tries to move away | 1 |
| | | | Aggressive reaction | 0 |
| | | Repeat to test habituation (10 times max, stop if dog is too anxious or uncomfortable) | Presence of habituation:[b] | |
| | | | Yes | 1 |
| | | | No | 0 |
| | 18 – Reactivity to visual stimuli | With the dog on a leash, approximately 3 m from the dog, without establishing visual contact, | Curious, wants to explore stimulus | 3 |
| | | | Does not show any reaction | 2 |
| | | | Fearful, tries to move away | 1 |

| | | | | |
|---|---|---|---|---|
| | | the experimenter opens an umbrella in a repeating action | Aggressive reaction | 0 |
| | | Repeat to test habituation (10 times, stop if dog is too anxious or uncomfortable) | Presence of habituation:[b] | |
| | | | Yes | 1 |
| | | | No | 0 |
| Behavior on Leash | 19 – Return to kennel | Return to the kennel on the leash. | Enters the kennel immediately | 2 |
| | | | Shows reluctance to enter | 1 |
| | | | Refuses to enter the kennel | 0 |

[a] Not applicable if the dog does not play with the ball (preceding question)
[b] Not applicable if the dog does not show any reaction to stimulus in the preceding question

**Appendix 5 – Ethogram for Stress-Related Behaviors**

The ethogram was adapted from a previous study[5]. The behaviors were live-scored during the training sessions at the prisons by two experimenters (ACVC and PB). Each experimenter tracked three dogs for each prison-batch combination, randomly attributed before the first session at the prisons. Each dog was scored during 5-minutes per training session, using a continuous sampling technique. The order in which the dogs were observed was kept constant across training sessions.

| Behavior | Definition |
|---|---|
| **Body turn** | Dog rotates its body (or head only) to the side, away from handler, in an attempt to avoid him/her, following an action such as looking at, approaching, or talking to the dog. Dog is in a tense or low posture. Ears are usually back. Tail can be down. Can be accompanied by lip licking or paw lift. |
| **Move away** | Dog takes one or a few steps away from handler (can be with rear or hind paws only), in an attempt to avoid or escape, following an action such as looking at, approaching, or talking to the dog. Dog is in a tense or low posture. Ears are usually back. Tail can be down. Can be accompanied by lip licking or paw lift. |
| **Crouch** | Dog lowers body (or head only) towards floor, usually lowering its head relative to torso (can be accompanied by blinking and dog's head can generally be turned away), bending legs and arching its back, following an action of handler, such as looking at, approaching, or talking to the dog. Ears are usually back. Tail can be down. Can be accompanied by lip licking or paw lift. |
| **Body shake** | Vigorous movement of the whole body side to side. |
| **Yawn** | Mouth opened wide briefly, then closed (may not close completely). |
| **Lip Lick** | Moving tongue around outside the mouth, touching other parts of the dog's face. Excluding times when the dog is focused on food/treat, up to 3 seconds after consuming food/treat. |

[5] Vieira de Castro AC, Fuchs D, Munhoz Morello G, Pastur S, de Sousa L, Olsson IAS. Does training method matter? Evidence for the negative impact of aversive-based methods on companion dog welfare. PloS One. 2020; 15(12): e0225023

**Appendix 6 – Ethogram for Overall Behavior State**

The ethogram was adapted from a previous study[5]. The behaviors were live-scored during the training sessions at the prisons by two experimenters (ACVC and PB). Each experimenter tracked three dogs for each prison-batch combination, randomly attributed before the first session at the prisons. Each dog was scored during 5-minutes per training session, using a scan-sampling 60s technique. The order in which the dogs were observed was kept constant across training sessions.

| Behavior State | Definition |
|---|---|
| **Tense** | Dog shows a combination of:<br>○ horizontal and tense body<br>○ tense muzzle<br>○ ears forward or back<br>○ tail held stiffly (high, neutral, or low) and still (in some cases it can be wagging)<br>○ lip licking<br>○ panting<br>○ paw lift<br>○ yawning<br>○ blinking |
| **Low** | Dog shows a combination of:<br>○ curved body<br>○ bent legs<br>○ low head<br>○ ears back<br>○ tail low, still or wagging<br>○ paw lift<br>○ lip licking<br>○ panting<br>○ yawning |
| **Relaxed** | Dog shows a combination of:<br>○ horizontal and relaxed body<br>○ ears in the normal position for dog<br>○ tail neutral, still or wagging slightly<br>○ can be panting |
| **Excited** | Dog shows a combination of:<br>○ horizontal body<br>○ ears forward or back<br>○ tail high or neutral, still or wagging<br>○ rapid or jerky movement<br>○ jumping<br>○ panting<br>○ play signals<br>○ body shake<br>○ may also yawn |
| **Unknown** | Dog is not visible, video recording is unclear, or the behavior of the dog cannot be clearly interpreted. |

**Appendix 7A – Demographic data for the TT and BET scorers**

| Name | Batches analyzed | Source | Type | Gender | Age range | Dog experience (0-10)* | Animal behavior experience (0-7)* |
|---|---|---|---|---|---|---|---|
| VS1 | 1, 2 | Video | Blind | F | 31-40 | 3 | 3 |
| VS2 | 1, 2 | Video | Blind | F | 18-30 | 4 | 1 |
| VS3 | 1, 2 | Video | Blind | M | 18-30 | 7 | 2 |
| VS4 | 3, 4 | Video | Blind | M | 18-30 | 2 | 2 |
| VS5 | 3 | Video | Blind | F | 18-30 | 1 | 1 |
| VS6 | 3 | Video | Blind | F | 18-30 | 7 | 4 |
| VS7 | 4 | Video | Blind | F | 18-30 | 4 | 2 |
| VS8 | 4 | Video | Blind | F | 18-30 | 0 | 2 |
| VS9 | 1, 2, 3, 4 | Video | Non-Blind | F | 18-30 | 5 | 4 |
| MQ | 1, 2, 3, 4 | Live | Non-Blind | F | 41-50 | 10 | 5 |
| ACVC | 1,2, 3, 4 | Live | Non-Blind | F | 31-40 | 10 | 7 |

*The scoring systems in Appendix 8b were used for rating.

**Appendix 7B – Dog Experience and Animal Behavior Experience Scoring Systems**

The scoring system was developed by the research team and then filled by the video and live scorers. Each scorer could choose multiple options for both questions and a composite score was assigned based on their responses.

| **Experience Level - Dogs** | **Score assigned** |
|---|---|
| None | 0 |
| Theoretical knowledge | 1 |
| Grown up with 1 or more dogs | 1 |
| Primary caregiver for a dog | 2 |
| Volunteered/worked with dogs in a shelter/clinic | 3 |
| Worked as a dog trainer or behaviorist | 3 |
| **Highest possible score** | **10** |

| **Experience Level - Animal Behavior** | **Score assigned** |
|---|---|
| None | 0 |
| Theoretical knowledge | 1 |
| Taken classes on animal behavior | 1 |
| Conducted/participated in studies that include observation of animal behavior (excluding dogs) | 2 |
| Conducted/participated in studies that include observation of dog behavior | 3 |
| **Highest possible score** | **7** |

**Appendix 8A – Inter-observer agreement results (TT)**

| | Batch 1 + Batch 2 | | Batch 3 | | Batch 4 | |
|---|---|---|---|---|---|---|
| **Item** | **ICC** | **p-value** | **ICC** | **p-value** | **ICC** | **p-value** |
| 1 | 0.875 | <0.001 | 0.792 | <0.001 | 0.894 | <0.001 |
| 2 | 0.662 | <0.001 | 0.767 | <0.001 | 0.641 | <0.001 |
| 3 | 0.821 | <0.001 | 0.942 | <0.001 | 0.583 | <0.001 |
| 4 | 0.902 | <0.001 | 0.948 | <0.001 | 0.550 | 0.001 |
| 5 | 0.862 | <0.001 | 0.955 | <0.001 | 0.666 | <0.001 |
| 6 | 0.721 | <0.001 | 0.976 | <0.001 | 0.794 | <0.001 |
| 7 | 0.884 | <0.001 | 0.964 | <0.001 | 0.570 | <0.001 |
| 8 | 0.594 | <0.001 | 0.866 | <0.001 | 0.900 | <0.001 |
| 9 | 0.594 | <0.001 | 0.891 | <0.001 | 0.858 | <0.001 |
| 10a | 0.567 | 0.002 | 0.905 | <0.001 | 0.620 | <0.001 |
| 10b | 0.647 | <0.001 | 0.841 | <0.001 | 0.768 | <0.001 |
| 10c | 0.765 | <0.001 | 0.720 | <0.001 | 0.910 | <0.001 |
| 11 | 0.860 | <0.001 | 0.974 | <0.001 | 0.949 | <0.001 |
| 12 | 0.760 | <0.001 | 0.890 | <0.001 | 0.901 | <0.001 |
| 13 | 0.532 | 0.004 | 0.947 | <0.001 | 0.853 | <0.001 |
| 14a | 0.777 | <0.001 | 0.986 | <0.001 | 0.990 | <0.001 |
| 14b | * | | 0.573 | 0.033 | 0.883 | <0.001 |
| 15 | 0.739 | <0.001 | 0.883 | <0.001 | 0.781 | <0.001 |
| 16a | 0.790 | <0.001 | 0.805 | <0.001 | 0.908 | <0.001 |
| 16b | 0.803 | <0.001 | 0.825 | <0.001 | 0.902 | <0.001 |
| 17a | 0.763 | <0.001 | 0.825 | <0.001 | 0.801 | <0.001 |
| 17b | 0.477 | 0.040 | 0.738 | <0.001 | 0.709 | <0.001 |
| 18a | 0.844 | <0.001 | 0.892 | <0.001 | 0.832 | <0.001 |
| 18b | 0.876 | <0.001 | 0.919 | <0.001 | 1.000 | ** |
| TT19 | 0.551 | <0.001 | 0.816 | <0.001 | 0.892 | <0.001 |

*n=0; analysis not performed

**no variability; all observers gave the same score

**Appendix 8B – Inter-observer agreement results (BET)**

| | Batch 1 + Batch 2 | | Batch 3 | | Batch 4 | |
|---|---|---|---|---|---|---|
| **Item** | **ICC** | **p-value** | **ICC** | **p-value** | **ICC** | **p-value** |
| 1 | 0.687 | <0.001 | 0.892 | <0.001 | 0.492 | 0.004 |
| 2 | 0.897 | <0.001 | 0.946 | <0.001 | 0.905 | <0.001 |
| 3 | 0.722 | <0.001 | 0.781 | <0.001 | 0.715 | <0.001 |
| 4 | 0.780 | <0.001 | 0.832 | <0.001 | 0.780 | <0.001 |
| 5 | 0.956 | <0.001 | 0.987 | <0.001 | 0.911 | <0.001 |
| 6 | 0.957 | <0.001 | 0.986 | <0.001 | 0.969 | <0.001 |
| 7 | 0.821 | <0.001 | 0.853 | <0.001 | 0.695 | <0.001 |

**Appendix 9 – Cognitive Bias Test Protocol**

The protocol was adapted from previous studies[5],[6]. The test was conducted in the same indoor room within the shelter (PATA) premises building where BETs and TTs were conducted. Two experimenters conducted the tests.

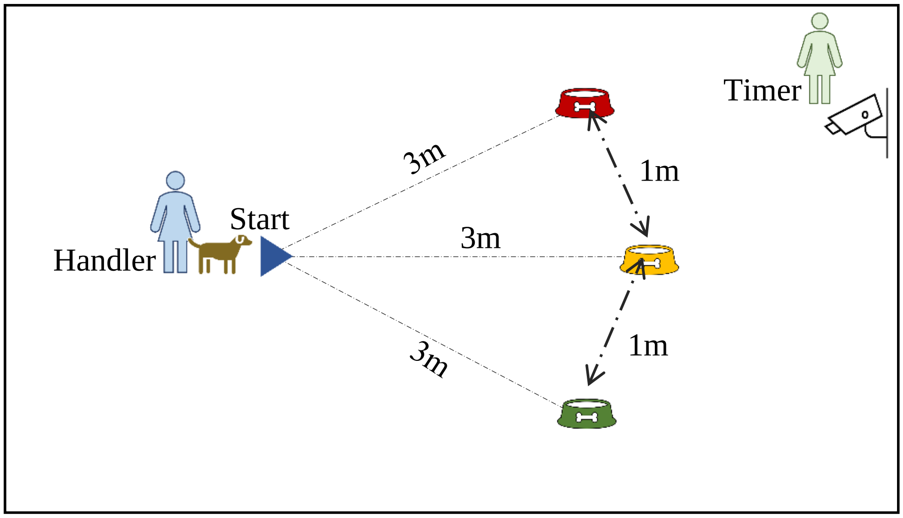


**Stage 1: Familiarization**

Prior to the start of the cognitive bias test, the dogs were given the opportunity to familiarize with the test room and the researchers. This consisted of a 3-minute period during which the dog was allowed to freely explore the room and engage with the researchers and the environment.

**Stage 2: Training**

The dog was on a slip-leash at all times. One researcher (ACVC or a MSc student) played the 'handler' and waited with the dog at the start point. The other researcher (PB), the 'timer', prepared the food bowl (with or without food) at the other end of the room, while giving their back to the dog. The food reward used was a piece of sausage of roughly 1.5 to 2 grams. One side of the room (from the dog's end) was set to be the 'positive' (P) location (the one where the bowl would be laid down containing food), and the other side the 'negative' (N) location (the one where the bowl would be laid down empty). To ensure that the dog and the 'handler' were blind to whether or not the bowl contained food during each trial, the bowl was baited out of their sight. Additionally, the food reward was rubbed onto the food bowl before every trial to prevent the influence of olfactory cues. The height of the food bowl was such that the dog was not able to visually judge the presence or absence of food from the start position.

After baiting (or not baiting) the bowl, the 'timer' placed it at one of the two training locations (P or N). The 'timer' then determined the start of the trial, by verbally signaling to the 'handler', upon which the 'handler' would release the dog at the start position. The 'handler' always led the dog to the start position on her right side. For dogs that had some difficulty noticing the bowl at the end of the room and would not leave the start position, during the first four trials the 'handler' would take one step towards the center of the two locations to encourage the dog to move. If the dog still did not move, the 'handler' lead the dog to the bowl or pointed it out. For the remaining trials, the 'handler' simply walked the dog to the start position and released him. After the dog reached the food bowl and (when applicable) ate the reward, the 'handler' would collect him and lead him back to start the next trial. The

[6] Duranton C, Horowitz A. Let me sniff! Nosework induces positive judgment bias in pet dogs. Applied Animal Behavior Science. 2019; 211, 61-66.

latency to reach the bowl, defined as the time elapsed between release at the start position and the dog putting its head in line with the edge of the bowl, was be recorded for each trial by the 'timer' using a stopwatch.

The position of the P and N locations was counterbalanced such that for half of the dogs, P was on the right side as they faced the test area, and for the other half it was on the left. Initially, each dog received two consecutive P trials followed by two N trials. Subsequently, P and N trials were presented in a pseudorandom order, with no more than two trials of the same type being presented consecutively.

All dogs will receive a minimum of 15 training trials to learn the discrimination between bowl locations. Dogs were considered to have learned an association between bowl location and food (the learning criterion) when, after a minimum of 15 trials, the longest latency to reach the P location was shorter than any of the latencies to reach the N location for the preceding three P and three N trials. Each trial lasted a maximum of 20 seconds. If the dog did not reach the bowl by that time, the trial was automatically terminated and a latency of 20 seconds was recorded.

**Stage 3: Testing**

Testing began once the learning criterion was achieved. Test trials were identical to training trials except that the bowl (empty) was placed at an ambiguous location, equally spaced along an arc 3 m from the dog's start position, between the P and N locations for the first test trial. For the pre-program assessment, the remaining (six) test trials were either P or N trials, presented in a pseudorandom order, with no more than two trials of the same type being presented consecutively. These trials were conducted to minimize learning effects to the post-program assessment.

**Appendix 10 – Number of dogs (N) for each of the paired t-tests performed for baseline cortisol analysis.**

| Comparison | N |
|---|---|
| B vs. C1 | 11 |
| B vs. C2 | 12 |
| B vs. C3 | 11 |
| B vs. F | 10 |
